# Gait asymmetry, and bilateral coordination of gait during a six-minute walk test in persons with multiple sclerosis

**DOI:** 10.1101/2020.05.13.093161

**Authors:** Meir Plotnik, Joanne M. Wagner, Gautam Adusumilli, Amihai Gottlieb, Robert T. Naismith

**Affiliations:** Center of Advanced Technologies in Rehabilitation, Sheba Medical Center, Ramat Gan, Israel; Department of Physiology and Pharmacology, Sackler Faculty of Medicine, Tel-Aviv University, Tel Aviv, Israel; Sagol School of Neuroscience, Tel Aviv University, Tel Aviv, Israel; Department of Physical Therapy and Athletic Training, Saint Louis University; Department of Neurology, Washington University in St. Louis

**Keywords:** Multiple sclerosis, Gait asymmetry, Bilateral coordination of gait, Six-minute walk test

## Abstract

Gait impairments in persons with multiple sclerosis (pwMS) underlying reduced walking endurance are still poorly understood. Thus, our objective was to assessed gait asymmetry (GA) and bilateral coordination of gait (BCG), among pwMS during the six-minute walk test (6MWT) and their association with disease severity. For this aim, we recruited ninety-two pwMS (age: 46.6 ± 7.9; 83% females) with a broad range of clinical disability who completed the 6MWT wearing gait analysis system. GA was assessed by comparing left and right swing times, and BCG by using the phase coordination index (PCI). Several functional and subjective gait assessments were performed. Results show that gait is more asymmetric and less coordinated as the disease progresses (p<.0001). Participants with mild MS showed significant better BCG as reflected by lower PCI values in comparison to the other two MS severity groups (severe: p =.001, moderate: p=.02). GA and PCI also deteriorated significantly with time during the 6MWT (p<.0001). GA and PCI (i.e., BCG) show somewhat weaker associations with clinical MS status than associations observed between functional and subjective gait assessments and MS status. Similar to other neurological cohorts, GA and PCI are important parameters to assess and to target in interventions among pwMS.

## Background

Multiple sclerosis (MS) is a degenerative, progressive, autoimmune disease of the central nervous system, often resulting in a continuous deterioration of walking ^1^. Hence, gait parameters, e.g. cadence, step-length, step-time, are impaired as compared to those measured in abled bodied individuals ^1–4^. This gait deterioration has been demonstrated as a decline in the ability to walk long distances, based on the 6-minute or 500-meter walk tests ^5,6^. One critical component of walking impairment is gait variability. Gait variability tends to change throughout the MS disease course, with greater variability in the higher levels of disability ^7,8^. Furthermore, gait variability is associated with increased fall risk ^9^.

### Potential association between MS pathology and difficulties in gait symmetry and bilateral coordination of gait (BCG)

Human gait requires a high degree of symmetry and coordination. Gait asymmetry is associated with reduced walking velocity ^10–12^ and increased energy expenditure ^13^. Spinal cord injury is frequent in persons with multiple sclerosis (pwMS), noted in 83% by MRI and up to 99% at autopsy ^14,15^. Spinal cord injury is associated with lower extremity sensorimotor deficits and impaired ambulation. It was previously reported that walking velocity in pwMS was reduced when in vivo diffusion tensor imaging (DTI) of the cervical spinal cord reveals myelin and tissue injury within posterior columns (PC) and lateral corticospinal tracts (CST) ^16^. Since CST injury in pwMS is asymmetric ^17^, we hypothesize that MS will be associated with increased gait asymmetry, since asymmetric lesions in the spinal white matter lesions correspond to asymmetric motor function ^18^.

Gait coordination is the ability to maintain a consistent phase-dependent cyclical relationship between different body segments or joints in both spatial and temporal domains ^19^. In humans, the control of right-left stepping, or bilateral coordination of gait (BCG), is hypothesized to be mediated by central pattern generators (CPGs) within lumbrosacral spinal locomotor centers ^13^. CPGs are local neuronal circuits that generate rhythmic stepping movements by alternating activity between groups of flexor and extensor muscles. CPGs are modulated by large fiber sensory afferents, along with supraspinal input ^20^. Damage within the spinal cord may limit afferent (PC) and efferent (CST) input to the CPGs and thus, contributes to impaired BCG ^21,22^. Thus, it is also hypothesized that MS is characterized by impaired BCG.

In the present study we assess GA by the evaluating how leg movements differ while walking (comparing swing times between the legs and symmetry of swing duration) ^23^. Left-right stepping coordination is assessed by the phase coordination index (PCI), a measure of the accuracy and consistency of the phase relationship between the step timing of the left and right legs ^24^. Only three reports directly assessed PCI or GA among pwMS. Gianfrancesco et al. compared between cane users (n=6) with non-users (n=5) and attributed lower values of PCI and GA (i.e., better coordination and lower asymmetry) to the former group ^25^. Kasser et al. compared PCI values before and after an acute aerobic exercise in pwMS and reported no significant change ^26^. However, the relation to other clinical parameters, such as disease severity, was not assessed in these papers. Recently, Shema◽Shiratzky et al.^27^ measured GA among pwMS during the 6-minute walk test (6MWT) and reported its relation to disease severity, however, only mild and moderate pwMS were assessed and PCI values were not measured in their study.

Thus, gait difficulties in left-right stepping coordination and gait symmetry in PwMS, measured by temporal gait metrics (cycle to cycle performance), has not yet been fully described.

### Study rationale

Characterizing GA and coordination impairments in MS, and their relations with functional (and subjective) gait assessments and with disease severity, would provide a more comprehensive picture about gait impairments in this cohort. GA and coordination impairments may prove useful in defining treatment targets and efficacy assessments for improving gait in MS. Assessing the relation of these parameters with disease severity could shed light on the progression of the MS and its effect on gait and risk of falls.

The objective of this study was to characterize GA and BCG in pwMS, during a long-distance walk (i.e., the 6MWT) and to assess their relationship with various, established gait assessments and with the clinical state of pwMS. The study is driven by two hypotheses: ***(1)*** Human gait requires a high degree of symmetry and coordination, both of which are potentially altered in MS. ***(2)*** The type of gait patterns in pwMS will be correlated with the level of impairment on key MS clinical outcomes describing, e.g., walking endurance and self-reported walking limitations.

## Methods

### Participants

A total of 92 pwMS were recruited through the John L. Trotter MS Center at Washington University School of Medicine. This cross-sectional study was approved by the local Human Research Protection Office/Institutional Review Board of the Washington University in St. Louis. All participants signed an informed consent prior to entering the study and all methods were performed in accordance with the relevant guidelines and regulations. Inclusion criteria were: >18 years of age; relapsing-remitting MS (RRMS), secondary-progressive MS (SPMS), or primary-progressive MS (PPMS); evidence of cervical spinal cord disease by symptoms or signs (e.g. Lhermitte’s sign, upper extremity weakness, upper extremity sensory symptoms), T2 lesions on clinical cervical MRI, and >180 days post-relapse. Exclusion criteria: confounders which could affect ambulation outside the spinal cord (e.g. poor vision, clear brainstem or cerebellar signs with MRI correlates and moderate-severe overall cerebral disease burden (>2.5 cm T2 lesion burden) ^16^ pregnancy, LE orthopedic conditions that limit ambulation, morbid obesity, cardiac pacemaker, metallic implant, and claustrophobia. Participants were then grouped by disease severity, i.e. mild (EDSS 0-2.5, n=60), moderate (EDSS 3-4, n=26) and severe (EDSS 4.5-6.5, n=6).

### Experimental protocol

The participants wore a six Opal motion sensor-based gait analysis system (APDM®, Portland, Oregon, USA). The 6MWT ^5^ was performed along a 50 foot walkway (i.e., participants turned around cones at each end of the 50 foot walkway). Spatiotemporal parameters of gait were collected by sensors worn on the ankles, wrists, lower back and chest, and transmitted to a wireless access device for storage on a mobile computer. Additional assessments included the Timed 25-foot Walk Test (T25FWT) ^28^, self-perceived walking limitation (12-item MS walking scale (MSWS-12v2)) ^29^ and balance confidence (Activities-specific balance confidence scale (ABC) ^30^.

### Gait cycle timing related Outcome measures

Outcomes were calculated based on ‘heel strike’ and ‘toe off’ timing obtained from the straight-line walking segments (180° turns at end of walkway were omitted):

1. Gait variability [%]: was defined by Stride time variability ^7,23^: coefficient of variation (CV) of the mean value of the stride time (multiple by 100 to obtain percentile unit). We termed this parameter stride CV. Stride CV was calculated separately for the left and the right legs and since values of the right and left legs were highly correlated (r=0.989; p< 0.001), we present data only for the right leg.
2. GA[%]= 100*|*ln*(*R_SW/L_SW*)|, where R_SW and L_SW stand for the mean value of right and left leg swing time, respectively ^23^.
3. Phase coordination index (PCI; [%] – for the quantification of BCG): A full description and derivation of the PCI metric is detailed elsewhere ^23^. Briefly, PCI is a metric that combines the accuracy and consistency of stepping phases generation with respect to the value of 180°, which represents the ideal anti-phased left–right stepping. Lower PCI values (also transformed to percentile unit; %) reflect a more consistent and more accurate phase generation, while higher values indicate a more impaired BCG ^23,31,32^.

### Functional and objective outcome measures

a. Distance covered during the 6MWT (ft) (reflective of gait speed).
b. Time (sec’) to complete the T25FWT
c. Transformed Score on the MSWS-12v2 ^33^ (higher score reflect perception of being more limited).
d. Transformed Score on the ABC (higher scores reflect more confidence in preforming balance capabilities related activities)

Gait variably, GA, PCI and Distance covered were calculated and measured for each minute of the 6MWT trail.

### Statistical analysis

For Gait variability, GA, PCI and distance covered, we performed a Repeated measures ANOVA treating each 1-minute interval (‘time’ effect) as a within subject level during the 6MWT, and the three severity subgroups (‘group’ effect) as a between-subject factor. When violations of the assumption of sphericity were observed, Greenhouse-Geisser estimates were used to correct the degrees of freedom. The Bonferroni method was used to correct for the post-hoc comparisons.

One-way between-groups ANOVAs and Bonferroni’s post-hoc tests compared the functional and objective outcome measures (apart of distance covered) among the MS severity subgroups.

Demographic and clinical factors which differed significantly between these groups (i.e. Age, Disease duration) were added as covariates in both of these analyses. Effect sizes were reported as Eta squared and Cohen’s d.

Correlation analyses between distance covered, stride CV, GA and PCI and the functional and subjective gait assessments were performed (Pearson) for the entire cohort and within each severity group. For the correlation analysis, we used the distance covered, stride CV, GA and PCI values measures and calculated over the entire 6MWT.

## Results

Demographic and clinical data of the sample appear in Table 1. Of the 92 participants, 60 were classified as having mild disability, 26 as moderate disability, and 6 as severe disability. As expected, significant group differences were found for age (F_(2,89)_ = 10.8, p < .001, η^2^ = 0.19) and disease duration (F_(2,89)_ = 3.39, p < .03, η^2^= 0.07). Post-hoc comparisons indicate younger age and shorter disease duration associated with mild disability. In light of these group differences, in the following analyses, we entered age and disease duration as covariates.

**Table 1:** Demographic and clinical data of the cohort.

|  | **Total (n = 92)** | **Mild (n = 60)** | **Moderate (n = 26)** | **Severe (n = 6)** |
| --- | --- | --- | --- | --- |
| **Age in years, mean ± SD (range)** | 46.6 ± 10.9 (22-66) | 43.2 ± 10.4 (22-66) | 53.0 ± 8.1 (33-65) | 54.1 ± 11.2 (34-63) |
| **EDSS, median (range)** | 2.0 (2.0-6.5) | 2.0 (0-2.5) | 3.0 (3-4) | 5.5 (5.0-6.5) |
| **Female, n (%)** | 76 (83%) | 49 (82%) | 21 (81%) | 6 (100%) |
| **Race (Caucasian, African-American, Asian), n (%)** | 83/8/1 (90,9,1) | 53/6/1 (88,10,2) | 24/2/0 (92,8,0) | 6/0/0 (100,0,0) |
| **Type of MS (RR/SP/PP), n (%)** | 88/2/2 (96%, 2%, 2%) | 59/0/1 (98%, 0%, 2%) | 23/2/1 (88%, 8%, 4%) | 6/0/0 (100%, 0%, 0%) |
| **Years since diagnosis, mean ± SD (range)** | 10.7 ± 7.9 (1-37) | 9.2 ± 7.0 (1-30) | 13.8 ± 7.8 (2-31) | 12.3 ± 12.6 (2-37) |

EDSS: expanded disability status scale, RR: relapsing remitting, SP: secondary progressive, PP: Primary progressive.

### Group (MS severity) and time effects on distance covered, gait variability, coordination and asymmetry

Significant group differences were found for distance covered (F_(2,87)_ = 20.7, p < .0001, η^2^ = 0.32), gait variability (i.e., Stride CV, F_(2,86)_ = 13.9, p < .0001, η^2^ = 0.24), PCI (F_(2,86)_ = 9.3, p < .0001, η^2^ = 0.17), and GA (F_(2,86)_ = 12.0, p < .0001, η^2^= 0.21; see figure 1 for post-hoc group to group comparisons).

**Figure 1.**
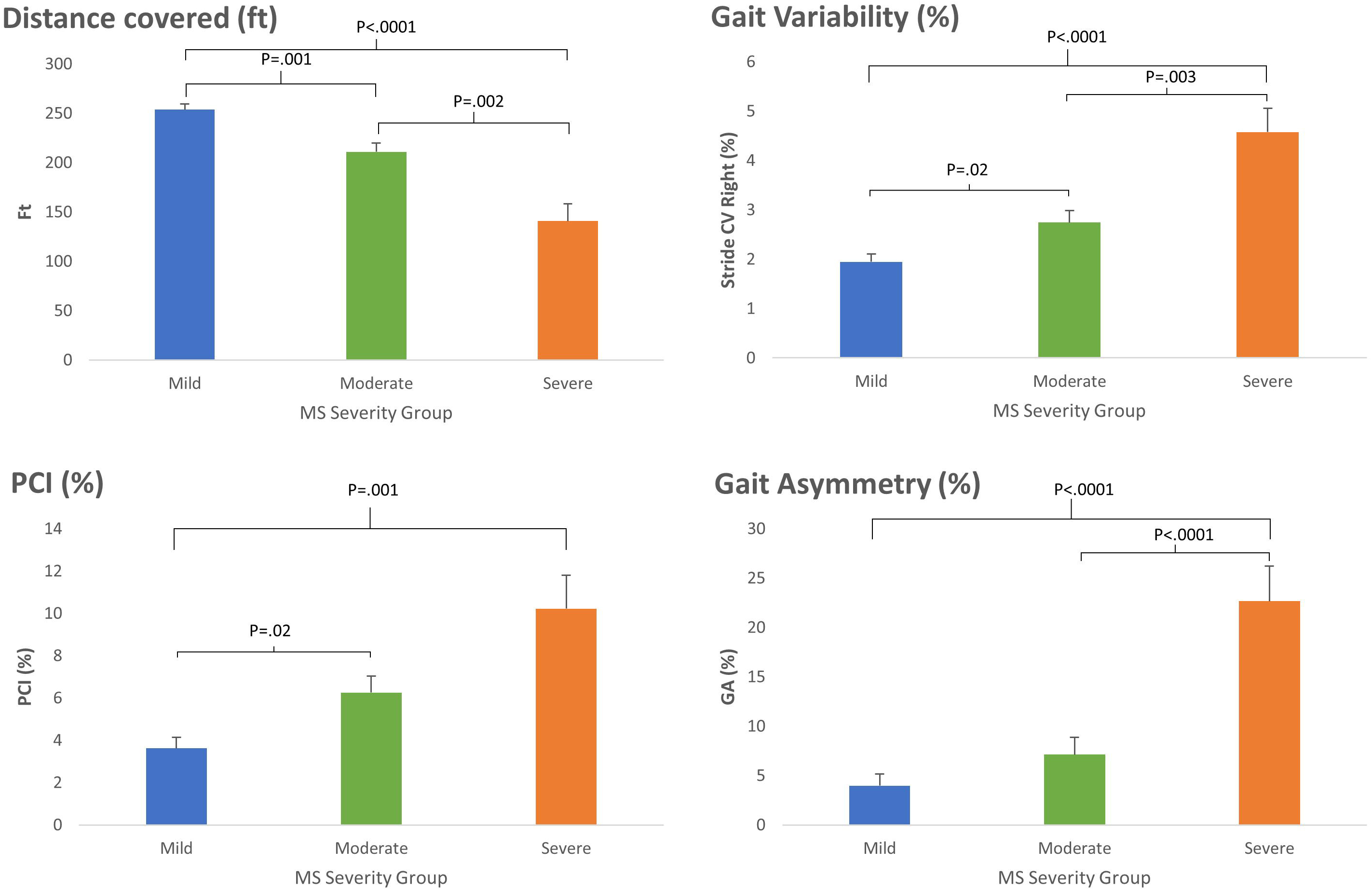
Mean values of distance covered, Stride time CV, GA and PCI during 6MWT in pwMS with different disease severity (see key). Bars represent mean values calculated based on the per-minute measures (see methods). Statistically significant differences in post-hoc group-to-group comparisons are marked above with horizontal brackets. Means and standard deviations are as followed: The severe disability group had greater asymmetry (GA: 22.6 ± 3.5) than the mild (GA: 3.9 ± 1.1) or moderate (GA: 7.1 ± 1.7) disability groups, and more impaired PCI (10.2 ± 1.5) than the mild disability group (3.6 ± 0.5). The moderate disability group had worse PCI (6.2 ± 0.5) compared to the mild disability group, and better PCI than the severe disability group. Gait Asymmetry did not significantly differ between the mild vs. moderate disability groups (p = 0.47). Distance covered significantly differed among all three MS severity groups (Severe: 141.6 ± 17.2, Moderate: 211.3 ± 8.3, Mild: 253.4 ± 5.4) and Gait Variability also significantly differed among all three MS severity groups (Severe: 4.5 ± 0.4, Moderate: 2.7 ± 0.2, Mild: 1.9 ± 0.1). Upper limit of the abscissa is 360 sec’.

Figure 2 depicts three examples of data sets from three participants, one from each severity level group. These examples are representative of the group differenced described above. Stride-to-stride time variability is larger for the more severely affected participants, as expressed by the larger range of values of the stride times in the participants with moderate and severe MS. Left-right stepping coordination (i.e., BCG) as expressed by the large variability in φ values (i.e., reduced consistency), and with values fairly ‘distant’ from the ‘ideal’ value of 180° is impaired in the participant with moderate MS (φ=173.0 ± 4.2°) as compared to the participant with mild MS (φ=179.4 ± 2.7°), and even worse in the participant with the severe MS (φ=187.3 ± 11.3°; see PCI values for all participants in figure 2). Similarly, the increased GA exhibited by the participants with moderate and severe MS is clearly demonstrated by the separation in the scatter of right (red) and left (green) swing times.

**Figure 2.**
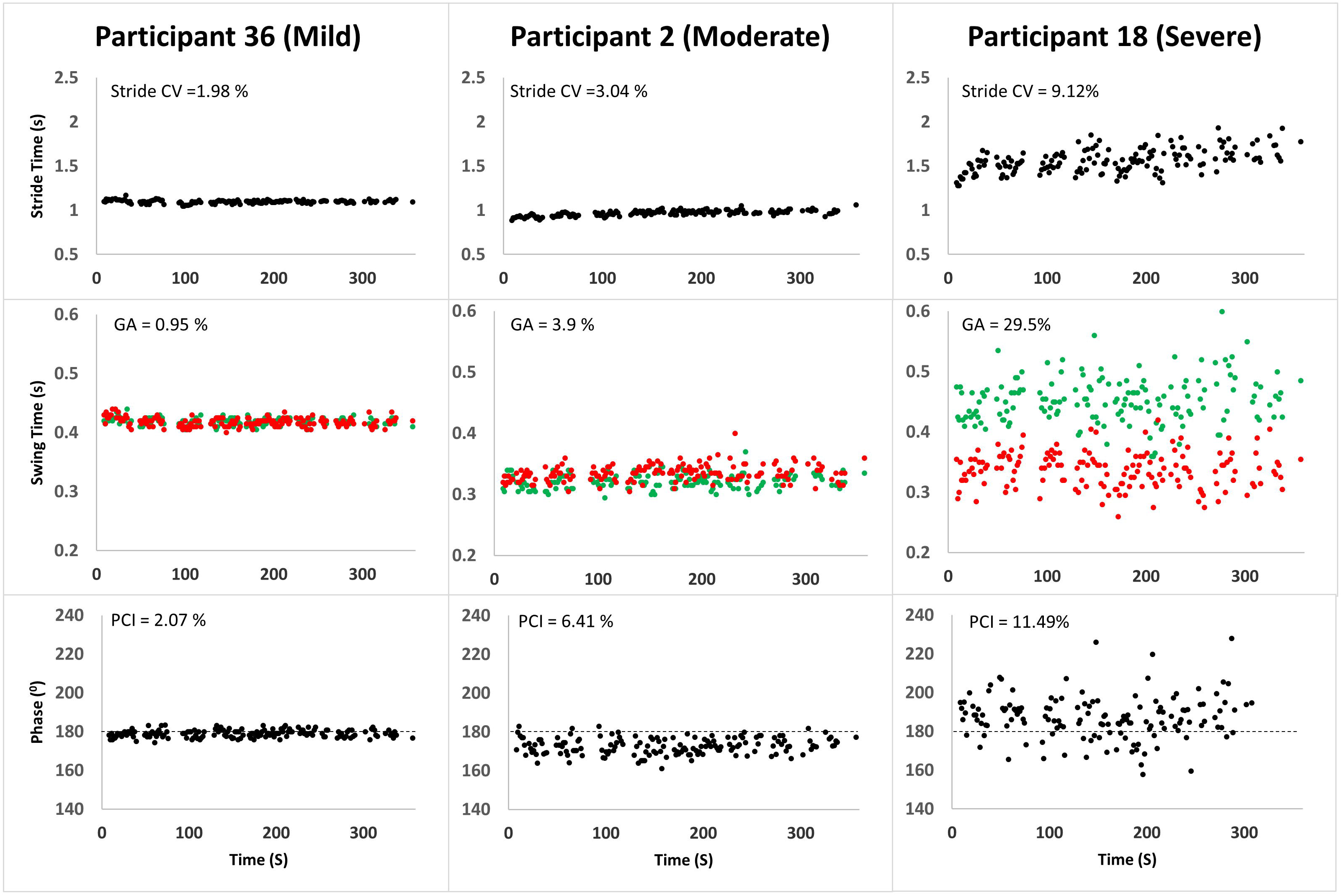
Three examples of data sets from three participants. Series of stride times (Top), left ‒right stepping phases (Middle), and swing times, (Bottom) are plotted for a 42 year old female patient with mild disease severity (left: EDSS = 2; 6MWT distance = 1514 ft; MSWWS-12v2 = 12; ABC = 98), a 59 year old female patient with moderate disease severity (middle: EDSS = 3.0, 6MWT distance = 1319 ft; MSWWS-12v2 = 38; ABC = 66), and a 48 year old female with severe disease severity (right: EDSS 6.5; 6MWT distance = 406 ft; MSWS12v2 = 76; ABC = 49). Red and green dots at the lower panels represent right and left swing times, respectively.

Significant time (i.e., potential fatigue) effects were found for PCI (F_(5,430)_ = 5.6, p < .0001, η^2^ = 0.06, due to a general decrease in coordination (i.e., larger PCI values) over time, in addition to a significant interaction between severity group and time, F_(10,430)_ = 4.9, p < .0001, η^2^ = 0.1, due to a larger increase in PCI values among the severe MS group. Further, significant time effects were also found for GA (F_(5,430)_ = 4.9, p < .0001, η^2^= 0.05, due to a general increase in asymmetry across groups, and in this case as well, a significant interaction was yielded between severity group and time, F_(10,430)_ = 5.3, p < .0001, η^2^ = 0.11, due to a larger increase in asymmetry among the severe MS group. Unexpectedly, time effect was not demonstrated with regards to distance covered, F_(5,435)_ = 0.7, p = .6, η^2^ = 0.008, or Gait variability (i.e., Stride CV, F_(5,430)_ = 1,8, p = .09, η^2^ = 0.02) nor interactions between Fatigue and Severity group regarding these measures.

### Functional and subjective gait assessments

Figure 3 depicts groups’ performance on the functional and subjective gait assessments. Eighty-one participants completed the T25FWT (52 form the mild group, 23 from the moderate group and six from the severe group). Group effect was statistically significant (F(_4,76_)=13.86, p <.001, η^2^ = 0.42), observing that participants with milder MS severity walk faster.

**Figure 3.**
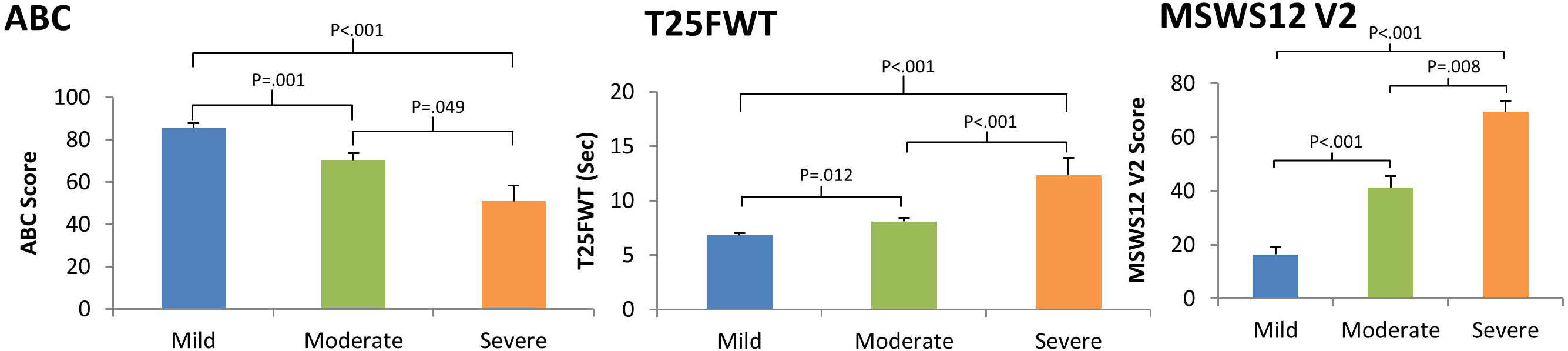
Mean values of ABC scores, MSWS12 V2 scores, and T25FWT seconds to complete in pwMS with different disease severity (see key). Statistically significant differences in post hoc group to group comparisons are marked above the horizontal brackets. Means, standard deviations and effect sizes are as followed: The severe disability group reported less balance confidence (ABC score: 50.9 ± 18.0), more self-perceived walking limitation (MSWS12 V2 score: 69.4 ± 9.9), and performed worse on the T25FWT seconds to complete (7.5 ± 2.2) than the moderate group (ABC score: 70.3 ± 16.3, d =1.12; MSWS12 V2 score: 41.2 ± 22.5, d =1.6; T25FWT seconds to complete: 12.3 ± 3.8, d =1.45) and compared to the mild group (ABC score: 85.4 ± 18.2, d =1.9; MSWS12 V2 score: 16.4 ± 20.4, d=3.29; T25FWT seconds to complete: 6.8 ± 1.4, d=1.9). Furthermore, the moderate severity group also reported less balance confidence, more self-perceived walking limitation, and performed worse on the T25FWT than the mild group (ABC, d=0.8; MSWS12 V2: d =1.14; T25FWT: d =0.8). Upper limit of the abscissa is 360 sec’.

Self-perceived walking limitations and balance confidence worsened with disease severity. Group effect was statistically significant for the MSWS-12v2 (F(_4,87_) = 14.48, p < .001, η^2^ = 0.40) and for the ABC test (F(_4,87_) = 7.49, p < .001, η^2^ = 0.25), as participants with milder MS severity reporting lower self-perceived gait limitations and higher balance confidence. Group to group post-hoc comparisons are detailed in figure 3.

### Correlation analyses

Statistically significant correlations were found between distance covered, PCI, GA and stride CV and clinical, functional and subjective outcomes (i.e., EDSS score, distance covered, MSWS12, T25WT and ABC) for the whole cohort (r_p_ = |0.29| – |0.60|, p < 0.01; see table 2 for details). PCI, GA and stride CV showed stronger inter correlations when evaluating data from pwMS participants with more severe disease (see table 3 for details). For example, PCI was strongly correlated with GA within the group with moderate MS (r_p=_ 0.86; p<0.01) and severe MS (r_p=_ 0.99; p<0.01), but not as strongly correlated within the group with mild MS (r_p=_ 0.36; p<0.01).

**Table 2:**
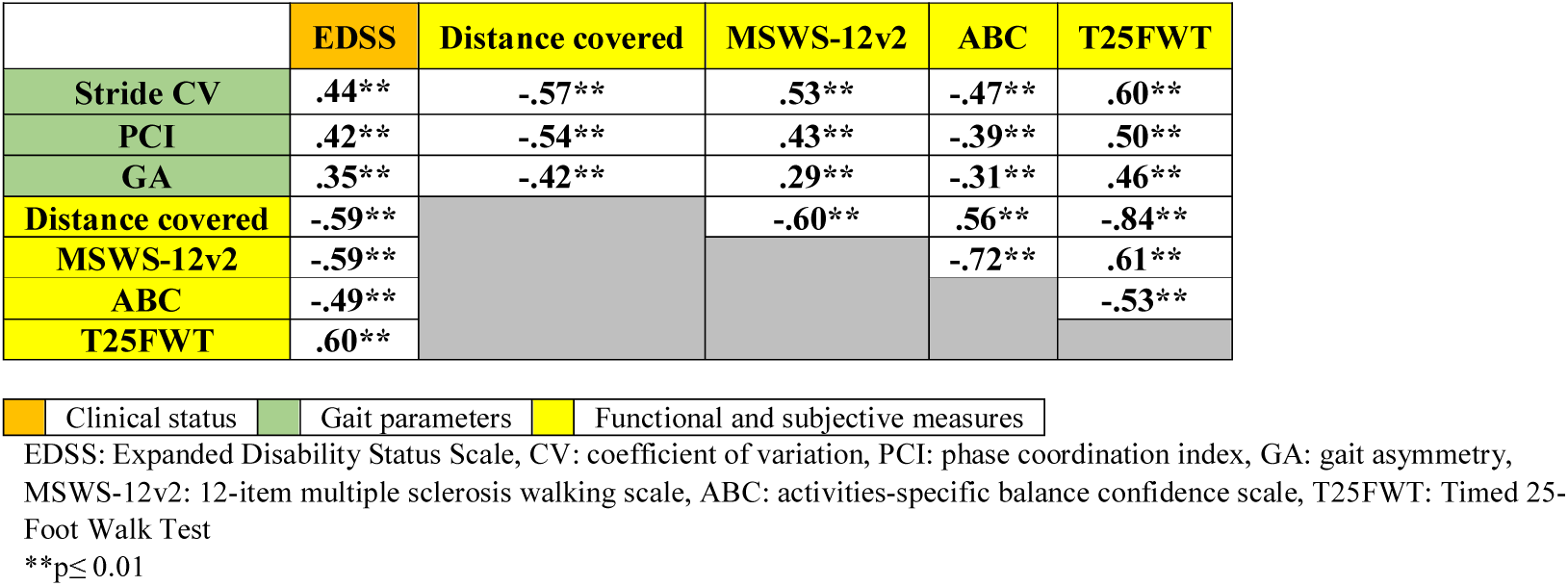
Pearson correlations between clinical status, gait parameters and functional and subjective gait assessments.

**Table 3:**
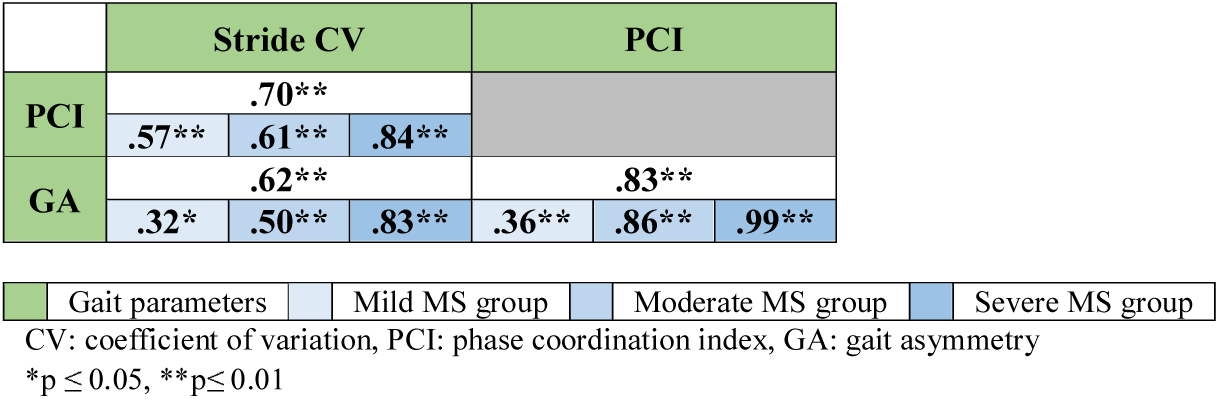
General and severity group Pearson correlations between the gait parameters.

## Discussion

### Summary of findings

This study which assessed gait asymmetry and impaired BCG among pwMS with spinal cord injury (i.e., excluding those with clear brainstem or cerebellar signs, see *Methods*) during relatively long-distance walk (i.e., 6MWT) observed a relationship with disease severity and functional and subjective gait assessments. In this cohort, gait was more asymmetric and less coordinated as the disease progressed. Participants with severe MS had larger GA values as compared to participants from the moderate and mild groups. Participants with mild MS showed significantly better BCG as reflected by lower PCI values in comparison to the other two MS severity groups. Furthermore, PCI and GA measured in participants with MS were highly inter-correlated, and correlated with gait variability, functional gait performance and subjective gait and balance assessments.

### Relation to previous findings

PCI and GA values in the current study found in the severe group were consistent with the PCI and GA values reported in Gianfrancesco et al. ^34^ measured among pwMS with severe MS (EDSS 4-5.5) while walking unassisted at a preferred walking speed. Despite the use of a different gait analysis system by these investigators utilized, GA and PCI appear to be robust measures characterizing the gait of pwMS.

Recently, Shema◽Shiratzky et al.^27^ also reported the results of a study in which various gait domains (e.g., pace, rhythm, variability, symmetry, and complexity) were measured and compared among mild- and moderate pwMS who performed the 6MWT while wearing body-fixed sensors. Though, they did not recruit participants with severe MS nor reported on PCI values. Their results show, similarly to the current findings, an association between gait deterioration, patient-reported gait disability and disease severity. However, in our study, symmetry had worsened during the 6MWT and variability did not while in Shema◽Shiratzky et al.’s study, variability had worsened, and symmetry did not change. Since we included participants with EDSS score lower than 2 and above 6, (not included by Shema- Shiratzky et al.) it might be the case that trends were missed or enhanced as a result of this truncation. Future studies may clarify this possibility.

While an able-bodied control group was not included in the present study, prior studies [32, 33] suggest gait parameters to be similar to our mild pwMS cohort. PCI values calculated using a similar gait analysis system were comparable (3.9 ± 13 %) to able-bodied controls with an age range similar to the age range of the mild pwMS cohort (45.4±3.6 y; n=20 vs 43.2±10.4; n=60, respectively). GA values were not reported on that study. Despite similar PCI values, the total distance covered by the present mild cohort (~1520 ft) was smaller than an able-bodied control group with similar age range (~2300 ft; ^35^). This suggests that PCI may not be the most sensitive or earliest parameters to coincide with walking impairment, and that deficits in walking endurance may precede impairment in the bilateral coordination of gait. In our study, pwMS with moderate disease severity exhibited PCI values of ~ 6.2% (fig. 1), larger than the PCI values observed in able-bodied participants from a comparable age group (PCI = 4.3%; ^36^). Thus, the PCI may differentiate pwMS from able-bodied persons as the disease progresses.

PCI values observed among participants from the severe MS group were similar to an elderly population. One study of able-bodied individuals in their eight decade of life demonstrated that their PCI was approximately 10%, quite comparable to the severe MS cohort PCI of 10.2% at a median age of 54 years ^37^. This observation illustrates the high degree of impaired coordination of gait in ambulatory pwMS with severe clinical disability. These data, taken together with a previous report of lower PCI and GA values (i.e., better coordination and lower asymmetry) in pwMS using a cane vs no cane ^25^, suggest the use of an assistive device may improve the BCG in similar pwMS.

PCI and GA were documented in other cohorts: stroke patients ^31^, persons with Parkinson’s disease (PD) ^23,38^ and elderly fallers ^38^. Similar to the results of the present study, PCI and GA were found to be associated with severity of the PD symptoms. Specifically, patients who suffer from the freezing of gait symptom exhibited higher values of GA ^39^ and PCI ^32^, indications for asymmetric gait, and impaired BCG.

Gait variability in pwMS has been assessed in many studies (see ^40^ for review). This gait parameter was found to be associated to disease severity, in agreement with the present results. It is worth noting that not all studies use the same parameter to describe gait variability. For instance, Kalron ^7^ and Socie et al. ^3^ used Step Length, Time and Width CV, Kaipust et al. ^41^ used Stride Length and Step Width CV.

Few studies have reported upon correlations between clinical, functional and subjective measures among pwMS. Goldman et al. ^5^ reported high correlation between the 6MWT and subjective quality of life and walking quality. Socie et al. ^3^ reported high correlations between disease severity and three measures of gait variability (e.g., r=0.51 between step length and EDSS values). Learmonth et al. ^42^ reported high correlations (r>|.62|) between EDSS and 6MWT distance, time to complete the T25FW and subjective walking quality. All these findings point to a positive relation between objective gait measures and subjective assessments of performance and quality of life.

### Relation between GA, BCG and MS

We found a strong correlation between GA and PCI (c.f., table 3). However, BCG differs from gait symmetry. BCG reflects the level of coordination between the ongoing stepping movements of both legs (i.e. the phase-dependent relationship between right and left heel strike over a number of steps) ^23,31^. Because people can have different swing durations for each leg with preservation of the phase relationship between the legs, an asymmetrical gait is not necessarily an uncoordinated gait ^23,43^. We propose that the PCI-GA correlation found in the present study may reflect the global impact of the pathogenicity on the ability to generate symmetric as well as coordinated gait.

The symmetrical gait commonly observed during normal walking can be attributed to the symmetric function of the central pattern generators (CPGs – the assumed neuronal substrate underlying rhythmical stepping movement of the lower limbs). Anti-phased stepping is an expression of the coordination between CPGs on both sides of the spine, most likely connected by commissural fibers ^21^. Damage to the CST in pwMS ^16^ most likely impact these neuronal substrates ^44^. Thus, we hypothesize that pwMS with asymmetrical CST are more likely to have an asymmetrical and less coordinated gait compared to pwMS with symmetrical CST, PC, or no injury. These hypotheses may be tested in future studies also involving imaging.

### Implications of findings, limitations, and future directions

We propose that stride CV, GA and PCI are important parameters to assess among pwMS, since they seem to deteriorate with disease progression. Stride CV, GA and PCI can be objectively assessed during home monitoring. While functional tests and subjective questionnaires require trained personnel, current light wearable sensors technologies (e.g. ^45,46^) allow an objective assessment of gait parameters like those at the focus of the present study. Longitudinal studies will be required to determine whether changes over time in these (and other gait parameters) have predictive diagnostic value regarding the progression of MS. Subtle impairments in gait performances have been demonstrated as prodromal signs for worsening due to PD ^36,47^. Further, gait coordination and gait asymmetry can be defined as targets for interventions aiming to improve gait performance ^48–50^. The efficacy of similar interventions in the context of fall prevention in MS ^51^ is still to be determined. Finally, our study is limited by the relatively small sample of pwMS with severe MS (n=6). Future studies should investigate a larger cohort of pwMS with severe disability.

In conclusion, GA and BCG are worse in pwMS who have more disability and disease progression, compared to those with mild to moderate disability. Longitudinal studies would help determine the rate of change in these parameters, and whether early changes have a predictive ability on disability and quality of life. Studies of physical therapy programs designed to target GA and BCG in pwMS may have potential to improve ambulation and falls.

## Acknowledgments

We thank Ms. Natalie Karlibach and Ms. Tamar Azrad from the Center of Advanced Technologies in Rehabilitation at Sheba Medical Center for technical support.

This research has been supported by NIH K12 HD055931 (Wagner, PI), and NMSS PP1940 (Wagner, PI). This publication was made possible by Grant Number UL1 RR024992 from the National Center for Research Resources (NCRR), a component of the National Institutes of Health (NIH) and NIH Roadmap for Medical Research. This research was also supported in part by NIH grants CO6 RR020092 and RR024992 (Washington University Institute of Clinical and Translational Sciences - Brain, Behavioral, and Performance Unit). Its contents are solely the responsibility of the authors and do not necessarily represent the official view of NCRR or NIH.

## Author Contributions

MP, JMW, GA, and RTN designed and performed the study. AG and MP did the data analysis, interpreted the data, prepared the figures and drafted the main manuscript text. JMW and RTN substantively revised the manuscript.

## Data availability

The datasets generated during and/or analysed during the current study are available from the corresponding author on reasonable request

## References

1 Motl, R. W. & Learmonth, Y. C. Neurological disability and its association with walking impairment in multiple sclerosis: brief review. Neurodegenerative disease management 4, 491–500 (2014).

2 Givon, U., Zeilig, G. & Achiron, A. Gait analysis in multiple sclerosis: characterization of temporal–spatial parameters using GAITRite functional ambulation system. Gait Posture 29, 138–142 (2009).

3 Socie, M. J., Motl, R. W., Pula, J. H., Sandroff, B. M. & Sosnoff, J. J. Gait variability and disability in multiple sclerosis. Gait Posture 38, 51–55 (2013).

4 Socie, M. J., Motl, R. W. & Sosnoff, J. J. Examination of spatiotemporal gait parameters during the 6-min walk in individuals with multiple sclerosis. Int J Rehabil Res 37, 311–316, doi:10.1097/MRR.0000000000000074 (2014).

5 Goldman, M. D., Marrie, R. A. & Cohen, J. A. Evaluation of the six-minute walk in multiple sclerosis subjects and healthy controls. Mult. Scler. 14, 383–390, doi:10.1177/1352458507082607 (2008).

6 Phan-Ba, R. et al. Motor fatigue measurement by distance-induced slow down of walking speed in multiple sclerosis. PLoS ONE 7, e34744, doi:10.1371/journal.pone.0034744 (2012).

7 Kalron, A. Gait variability across the disability spectrum in people with multiple sclerosis. Journal of the Neurological Sciences 361, 1–6, doi:10.1016/j.jns.2015.12.012 (2016).

8 Kalron, A. & Frid, L. The “butterfly diagram”: A gait marker for neurological and cerebellar impairment in people with multiple sclerosis. Journal of the Neurological Sciences 358, 92–100, doi:10.1016/j.jns.2015.08.028 (2015).

9 Kalron, A. Association between gait variability, falls and mobility in people with multiple sclerosis: A specific observation on the EDSS 4.0-4.5 level. NeuroRehabilitation 40, 579–585, doi:10.3233/NRE-171445 (2017).

10 Hsu, A.-L., Tang, P.-F. & Jan, M.-H. Analysis of impairments influencing gait velocity and asymmetry of hemiplegic patients after mild to moderate stroke. Arch Phys Med Rehabil 84, 1185–1193 (2003).

11 Lin, P.-Y., Yang, Y.-R., Cheng, S.-J. & Wang, R.-Y. The relation between ankle impairments and gait velocity and symmetry in people with stroke. Arch Phys Med Rehabil 87, 562–568, doi:10.1016/j.apmr.2005.12.042 (2006).

12 Patterson, K. K. et al. Gait asymmetry in community-ambulating stroke survivors. Arch Phys Med Rehabil 89, 304–310, doi:10.1016/j.apmr.2007.08.142 (2008).

13 Lewek, M. D., Osborn, A. J. & Wutzke, C. J. The influence of mechanically and physiologically imposed stiff-knee gait patterns on the energy cost of walking. Arch Phys Med Rehabil 93, 123–128, doi:10.1016/j.apmr.2011.08.019 (2012).

14 Bot, J. et al. Spinal cord abnormalities in recently diagnosed MS patients added value of spinal MRI examination. Neurology 62, 226–233 (2004).

15 Ikuta, F. & Zimmerman, H. Distribution of plaques in seventy autopsy cases of multiple sclerosis in the United States. Neurology 26, 26–28 (1976).

16 Naismith, R. T. et al. Spinal cord tract diffusion tensor imaging reveals disability substrate in demyelinating disease. Neurology 80, 2201–2209, doi:10.1212/WNL.0b013e318296e8f1 (2013).

17 Reich, D. S. et al. Multiparametric magnetic resonance imaging analysis of the corticospinal tract in multiple sclerosis. Neuroimage 38, 271–279 (2007).

18 von Meyenburg, J. et al. Spinal cord diffusion-tensor imaging and motor-evoked potentials in multiple sclerosis patients: microstructural and functional asymmetry. Radiology 267, 869–879 (2013).

19 Krasovsky, T. & Levin, M. F. Review: toward a better understanding of coordination in healthy and poststroke gait. Neurorehabil Neural Repair 24, 213–224, doi:10.1177/1545968309348509 (2010).

20 Dietz, V. Spinal cord pattern generators for locomotion. Clin Neurophysiol 114, 1379–1389 (2003).

21 Guertin, P. A. Central pattern generator for locomotion: anatomical, physiological, and pathophysiological considerations. Frontiers in neurology 3, 183 (2013).

22 OConnor, S. M. & Donelan, J. M. Fast visual prediction and slow optimization of preferred walking speed. American Journal of Physiology-Heart and Circulatory Physiology (2012).

23 Plotnik, M., Giladi, N. & Hausdorff, J. M. A new measure for quantifying the bilateral coordination of human gait: effects of aging and Parkinson’s disease. Exp Brain Res 181, 561–570, doi:10.1007/s00221-007-0955-7 (2007).

24 Plotnik, M., Giladi, N. & Hausdorff, J. M. A new measure for quantifying the bilateral coordination of human gait: effects of aging and Parkinson’s disease. Exp Brain Res 181, 561–570 (2007).

25 Gianfrancesco, M. A. et al. Speed- and cane-related alterations in gait parameters in individuals with multiple sclerosis. Gait Posture 33, 140–142, doi:10.1016/j.gaitpost.2010.09.016 (2011).

26 Kasser, S. L., Jacobs, J. V., Sibold, J., Marcus, A. & Cole, L. Employing Body-Worn Sensors to Detect Changes in Balance and Mobility After Acute Aerobic Exercise in Adults with Multiple Sclerosis. International Journal of MS Care (2019).

27 Shema-Shiratzky, S. et al. Deterioration of specific aspects of gait during the instrumented 6-min walk test among people with multiple sclerosis. Journal of neurology, 1–9 (2019).

28 Cutter, G. R. et al. Development of a multiple sclerosis functional composite as a clinical trial outcome measure. Brain 122 (Pt 5), 871–882 (1999).

29 Hobart, J. C., Riazi, A., Lamping, D. L., Fitzpatrick, R. & Thompson, A. J. Measuring the impact of MS on walking ability: the 12-Item MS Walking Scale (MSWS-12). Neurology 60, 31–36 (2003).

30 Powell, L. E. & Myers, A. M. The Activities-specific Balance Confidence (ABC) Scale. J. Gerontol. A Biol. Sci. Med. Sci. 50A, M28–34 (1995).

31 Meijer, R. et al. Markedly impaired bilateral coordination of gait in post-stroke patients: Is this deficit distinct from asymmetry? A cohort study. J Neuroeng Rehabil 8, 23, doi:10.1186/1743-0003-8-23 (2011).

32 Plotnik, M., Giladi, N. & Hausdorff, J. M. Bilateral coordination of walking and freezing of gait in Parkinson’s disease. European Journal of Neuroscience 27, 1999–2006 (2008).

33 Nilsagård, Y., Gunnarsson, L.-G. & Denison, E. Self-perceived limitations of gait in persons with multiple sclerosis. Advances in physiotherapy 9, 136–143 (2007).

34 Gianfrancesco, M. A. et al. Speed-and cane-related alterations in gait parameters in individuals with multiple sclerosis. Gait Posture 33, 140–142 (2011).

35 Pajek, M. B. et al. Six-minute walk test in renal failure patients: representative results, performance analysis and perceived dyspnea predictors. PLoS ONE 11, e0150414 (2016).

36 Mirelman, A. et al. Effects of aging on arm swing during gait: the role of gait speed and dual tasking. PLoS ONE 10, e0136043 (2015).

37 Gimmon, Y. et al. Gait Coordination Deteriorates in Independent Old-Old Adults. Journal of aging and physical activity 26, 382–389 (2018).

38 Yogev, G., Plotnik, M., Peretz, C., Giladi, N. & Hausdorff, J. M. Gait asymmetry in patients with Parkinson’s disease and elderly fallers: when does the bilateral coordination of gait require attention? Exp Brain Res 177, 336–346, doi:10.1007/s00221-006-0676-3 (2007).

39 Plotnik, M., Giladi, N., Balash, Y., Peretz, C. & Hausdorff, J. M. Is freezing of gait in Parkinson’s disease related to asymmetric motor function? Annals of Neurology: Official Journal of the American Neurological Association and the Child Neurology Society 57, 656–663 (2005).

40 Socie, M. J. & Sosnoff, J. J. Gait variability and multiple sclerosis. Multiple sclerosis international 2013 (2013).

41 Kaipust, J. P., Huisinga, J. M., Filipi, M. & Stergiou, N. Gait variability measures reveal differences between multiple sclerosis patients and healthy controls. Motor control 16, 229–244 (2012).

42 Learmonth, Y. C., Motl, R. W., Sandroff, B. M., Pula, J. H. & Cadavid, D. Validation of patient determined disease steps (PDDS) scale scores in persons with multiple sclerosis. BMC neurology 13, 37 (2013).

43 Plotnik, M. & Hausdorff, J. M. The role of gait rhythmicity and bilateral coordination of stepping in the pathophysiology of freezing of gait in Parkinson’s disease. Movement disorders: official journal of the Movement Disorder Society 23, S444–S450 (2008).

44 DeLuca, G., Ebers, G. & Esiri, M. Axonal loss in multiple sclerosis: a pathological survey of the corticospinal and sensory tracts. Brain 127, 1009–1018 (2004).

45 Newland, P. et al. Exploring the feasibility and acceptability of sensor monitoring of gait and falls in the homes of persons with multiple sclerosis. Gait Posture 49, 277–282 (2016).

46 Shammas, L. et al. Home-based system for physical activity monitoring in patients with multiple sclerosis (Pilot study). Biomedical engineering online 13, 10 (2014).

47 van den Heuvel, L. et al. Actigraphy detects greater intra-individual variability during gait in non-manifesting LRRK2 mutation carriers. Journal of Parkinson’s disease 8, 131–139 (2018).

48 Fasano, A. et al. Split-belt locomotion in Parkinson’s disease links asymmetry, dyscoordination and sequence effect. Gait Posture 48, 6–12 (2016).

49 Reisman, D. S., McLean, H., Keller, J., Danks, K. A. & Bastian, A. J. Repeated split-belt treadmill training improves poststroke step length asymmetry. Neurorehabil Neural Repair 27, 460–468 (2013).

50 Ricciardi, L. et al. Working on asymmetry in Parkinson’s disease: randomized, controlled pilot study. Neurological Sciences 36, 1337–1343 (2015).

51 Matsuda, P. N. et al. Falls in multiple sclerosis. PM&R 3, 624–632 (2011).

